# Accessing Mesoscale Functional Organization from Standard Functional MRI Leveraging AI-Based Super-Resolution

**DOI:** 10.64898/2026.11.07.765069

**Authors:** Shahin Nasr, Hongwei B. Li, Dina Zemlyanker, Laurentius Huber, Matthew S. Rosen, Juan E. Iglesias

## Abstract

Access to the fine-scale functional organization of the human brain is essential for understanding cognition and neurological disorders yet remains beyond the reach of conventional functional MRI (fMRI) because of its limited spatial resolution and contrast-to-noise ratio. To overcome this limitation, we developed **f-LENNS** (functional Localization Enhancement via Neural-Net-Sharpening), an artificial intelligence (AI)-based framework that enhances the effective spatial resolution of reconstructed fMRI magnitude images. The model was trained exclusively on high-resolution synthetic MRI data using extensive domain randomization and no adversarial or generative components. This approach enables robust out-of-the-box generalization (without retraining) to experimental datasets obtained with diverse acquisitions, while effectively curbing hallucination. Our experiments show that our publicly available model substantially improves the spatial fidelity of functional activation maps across multiple imaging paradigms and generalizes to a cohort of individuals with atypical brain function caused by amblyopia (lazy eye). F-LENNS consistently outperforms conventional resolution enhancement approaches, including interpolation-based upsampling and sharpening methods. These findings demonstrate that AI-driven super-resolution combined with domain randomization can overcome domain shifts arising from differences in MRI acquisition, enabling scalable, fine-scale functional brain mapping with widely available MRI hardware.

**Summary statement:** Precision functional magnetic resonance imaging (fMRI) at the mesoscale is transforming our understanding of human brain functions by enabling the visualization of fine-grained functional organization in the human brain. Recent advances have shown that precision fMRI can reveal heterogeneous cortical architecture, resolve mesoscale processing streams underlying sensory perception, attention, and decision-making, identify functional alterations associated with neurological disorders, and help explain inter-individual variability in cognition. However, the widespread adoption of these approaches remains constrained by the limited availability of ultra-high-field MRI systems.

Here, we introduce f-LENNS (functional Localization Enhancement via Neural-Net Sharpening), an artificial intelligence (AI)-based framework that broadens access to mesoscale functional neuroimaging by enhancing the effective spatial resolution of conventional fMRI. The model is trained exclusively on domain-randomized synthetic MRI data and generalizes to diverse datasets *without retraining*: f-LENNS can super-resolve data obtained with different acquisition parameters and at different magnetic field strengths (3T and 7T), while minimizing hallucination artifacts. F-LENNS substantially improves the detectability and spatial fidelity of fine-scale functional organization defined by stimulus-evoked activity and accurately recovers mesoscale functional patterns from conventional-resolution fMRI. Importantly, the same model generalizes to individuals with atypical brain function, revealing disorder-related functional abnormalities without data- or disease-specific retraining.

Together, these findings demonstrate that domain randomization enables AI-based super-resolution models to overcome domain shifts arising from heterogeneous MRI hardware and acquisition protocols. Compared with conventional approaches, including interpolation-based upsampling, AFNI sharpening, and AI models trained on real functional and/or structural MRI data, f-LENNS consistently provides superior reconstruction of fine-scale functional organization. By enabling precision functional imaging in individual participants using widely available MRI systems, f-LENNS expands access to mesoscale brain mapping and represents an important step toward scalable, individualized functional neuroimaging and precision medicine.

## 1. Introduction

High-resolution functional magnetic resonance imaging (fMRI) has enabled the mapping of mesoscale functional organization across the human brain, providing unprecedented insight into the neural circuits underlying perception^1–6^, cognition^7,8^, and neurological disorders^9–13^. However, achieving such spatial resolution typically requires ultra-high-field (UHF) MRI systems together with extensive signal averaging to compensate for the reduced signal- and contrast-to-noise ratios associated with smaller voxel sizes^14^. Given the limited availability of UHF scanners worldwide and practical constraints on scan duration, particularly in studies involving individuals with atypical brain function, the widespread adoption of high-resolution fMRI remains limited.

In the past, AI-based super-resolution (SR) methods, particularly those based on convolutional neural networks (CNNs)^15,16^ have successfully enhanced the spatial resolution of structural MRI acquired under conventional clinical protocols^17–20^. Their ability to recover realistic high-frequency image content has been further improved through adversarial learning (e.g., generative adversarial networks (GANs)^21–24^ and deep generative models (e.g., diffusion models)^25,26^), which greatly enhance the realism of the outputs^27,28^ – albeit at the expense of increasing potential hallucination artifacts. Despite these advances, extending AI-based SR to fMRI has proven considerably more challenging.

A central obstacle is the domain-shift problem: existing AI-based SR performance degrades sharply when imaging conditions differ from those seen during training. In fMRI, such shifts arise from variations in pulse sequence and contrast mechanism (e.g., gradient- or spin-echo Imaging^29^, and vascular-space-occupancy^30^), acquisition parameters (e.g., resolution and slice orientation), and field strength (3T vs. 7T). As a result, existing AI-based SR approaches for fMRI remain scarce and are typically restricted to either their usage for denoising at the level of image reconstruction^31,32^, or to subject-specific enhancement of data from a single site, protocol, and/or stimulus paradigm^33^.

Here, we introduce f-LENNS (functional Localization Enhancement via Neural Network Sharpening), an open-source AI framework that enables fMRI SR across diverse acquisition protocols. F-LENNS is specifically tailored to functional imaging through an fMRI-oriented training strategy that emphasizes the spatial resolutions commonly encountered in fMRI acquisitions. By learning to enhance spatial resolution while preserving the original image contrast, the model can be applied across diverse fMRI acquisition protocols without retraining.

F-LENNS combines a modern CNN architecture with domain randomization^34^ and extensive data augmentation^35^ to substantially mitigate domain shift (Fig. 1). While domain randomization effectively minimizes hallucination artifacts, we curb these even further by deliberately avoiding adversarial and generative components. Rather than attempting to exploit temporal information, f-LENNS processes each fMRI volume independently, thereby avoiding the introduction of temporal correlations that would be intractable and nonlinear, biasing downstream analyses. Furthermore, working one frame at a time makes f-LENNS compatible with any repetition time (TR).

**Figure 1.**
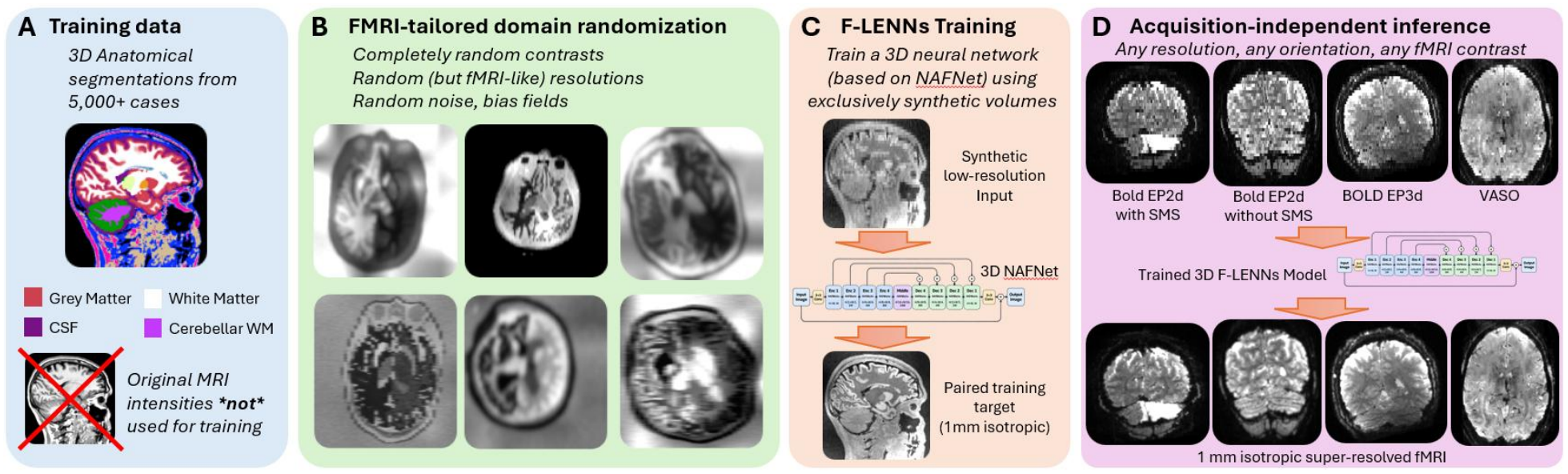
Overview of f-LENNS and its acquisition-independent SR framework. Anatomical segmentations from 5,241 individuals across 10 publicly available MRI datasets provide the anatomical information used to generate training data; the corresponding MRI intensity images are not used for training. Synthetic MRI volumes are generated from the anatomical segmentations using domain randomization to vary image contrast, spatial resolution, and acquisition characteristics. This procedure produces heterogeneous synthetic low-resolution inputs, which are resampled to a common 1-mm isotropic grid and paired with high-resolution target volumes. The network receives only the resampled image intensities, without explicit information about the original input resolution. Once trained, the same fixed model can be applied to real fMRI acquisitions with different contrasts and voxel sizes, including BOLD and VASO data, to generate 1-mm isotropic super-resolved fMRI without dataset-specific retraining.

As a result, f-LENNS can be applied directly (without retraining or fine-tuning) to unprocessed fMRI data acquired across different magnetic field strengths (3T and 7T), pulse sequences, spatial resolutions, and experimental paradigms. We demonstrate that f-LENNS substantially improves the spatial fidelity of functional activation maps while accurately recovering mesoscale functional organization across a broad range of datasets.

Importantly, unlike AI-based image-reconstruction methods integrated into the acquisition pipeline (e.g., Siemen’s proprietary Deep Resolve) and therefore requiring prospective implementation on compatible MRI systems, f-LENNS operates entirely as a post-processing framework on reconstructed magnitude images, without requiring complex-valued MRI data. Consequently, it can be applied retrospectively to both newly acquired and previously collected datasets (including large public neuroimaging repositories) without access to raw k-space data, scanner-specific reconstruction pipelines, retraining, or protocol-specific optimization. The same pretrained model works out of the box across heterogeneous datasets, enabling scalable access to precision functional neuroimaging using widely available MRI hardware.

## 2. Results

### 2.1. Overview of the f-LENNS framework

F-LENNS is a three-dimensional (3D) CNN designed for acquisition-independent SR of fMRI magnitude images. The network operates exclusively on real-valued image intensities, without using complex-valued MRI data or phase information. The model is based on NAFNet^36^, a lightweight image-restoration architecture that has shown strong performance across a range of 2D restoration tasks while maintaining low computational cost, making it well suited for extension to memory-intensive 3D fMRI data (Fig. 1). F-LENNS is trained exclusively on synthetic MRI volumes generated via Gaussian distributions conditioned on anatomical segmentations, rather than on real fMRI data. This strategy largely reduces hallucination at test time, while also enabling so-called “domain randomization”: During training, we introduce extensive variability in the contrast, resolution, and artifacts of synthetic images, while keeping the desired outputs at 1 mm isotropic resolution. Importantly, the distribution of input resolutions is specifically tailored to reflect those commonly encountered in fMRI acquisitions (i.e., roughly isotropic). The network learns to recover high-resolution image details while preserving the original image contrast, enabling its application across different fMRI acquisition protocols.

Importantly, all inputs are first resampled to a common 1 mm isotropic grid, and the network is not explicitly informed of their original voxel size. This has three beneficial effects: *(i)* f-LENNS super-resolves to 1 mm isotropic, irrespective of the resolution of the input; *(ii)* f-LENNS is compatible with inputs with heterogeneous contrasts, including blood-oxygen-level-dependent (BOLD) and vascular-space-occupancy (VASO) acquisitions; and *(iii)* f-LENNS works “out of the box” and does not require dataset-specific retraining. While domain randomization has a slightly lower performance ceiling than approaches based on real images when training and test data are exactly from the same distribution, they have been shown to outperform such supervised approaches (and also methods based on more accurate simulations), even when the domain shift is small^37^.

We evaluated f-LENNS by testing its ability to recover mesoscale functional organization in the human visual system across datasets acquired under diverse imaging conditions and in individuals with neurotypical and atypical visual function. As benchmarks, we additionally trained models directly on real resting-state fMRI from an independent cohort not included in the evaluation datasets. Architectural details, synthetic-data generation, training procedures, and benchmark-model construction are described in Methods.

### 2.2. F-LENNS improves the quality of raw fMRI images

To validate f-LENNS performance, the trained model was applied to fMRI data from individuals with neurotypical brain function. In contrast to the training data (i.e., synthetic datasets), the validation data was collected during various perceptual tasks. In this validation, the original high-resolution fMRI dataset, acquired at 1 mm isotropic resolution on a 7T scanner from eleven individuals, was first downsampled to either 2 mm or 3 mm isotropic resolution^38^ nd then super-resolved back to 1 mm using the trained model. While the lower resolution is synthetic, this approach enabled us to assess the performance of f-LENNS as a function of resolution independently of the test-retest uncertainty.

F-LENNS produced super-resolved images with significantly improved image fidelity compared with standard upsampling and AFNI sharpening approaches across different brain areas, including cortical, subcortical, and cerebellum regions (Fig. 2). These improvements were associated with reduced mean absolute error (L1) and mean squared error (L2) and increased structural similarity and peak signal-to-noise ratio (PSNR) relative to the upsampled and sharpened images (t(10)>11.18, *p*<10^-3^, corrected for multiple comparisons). These improvements were consistently observed for both 2 mm and 3 mm inputs.

**Figure 2.**
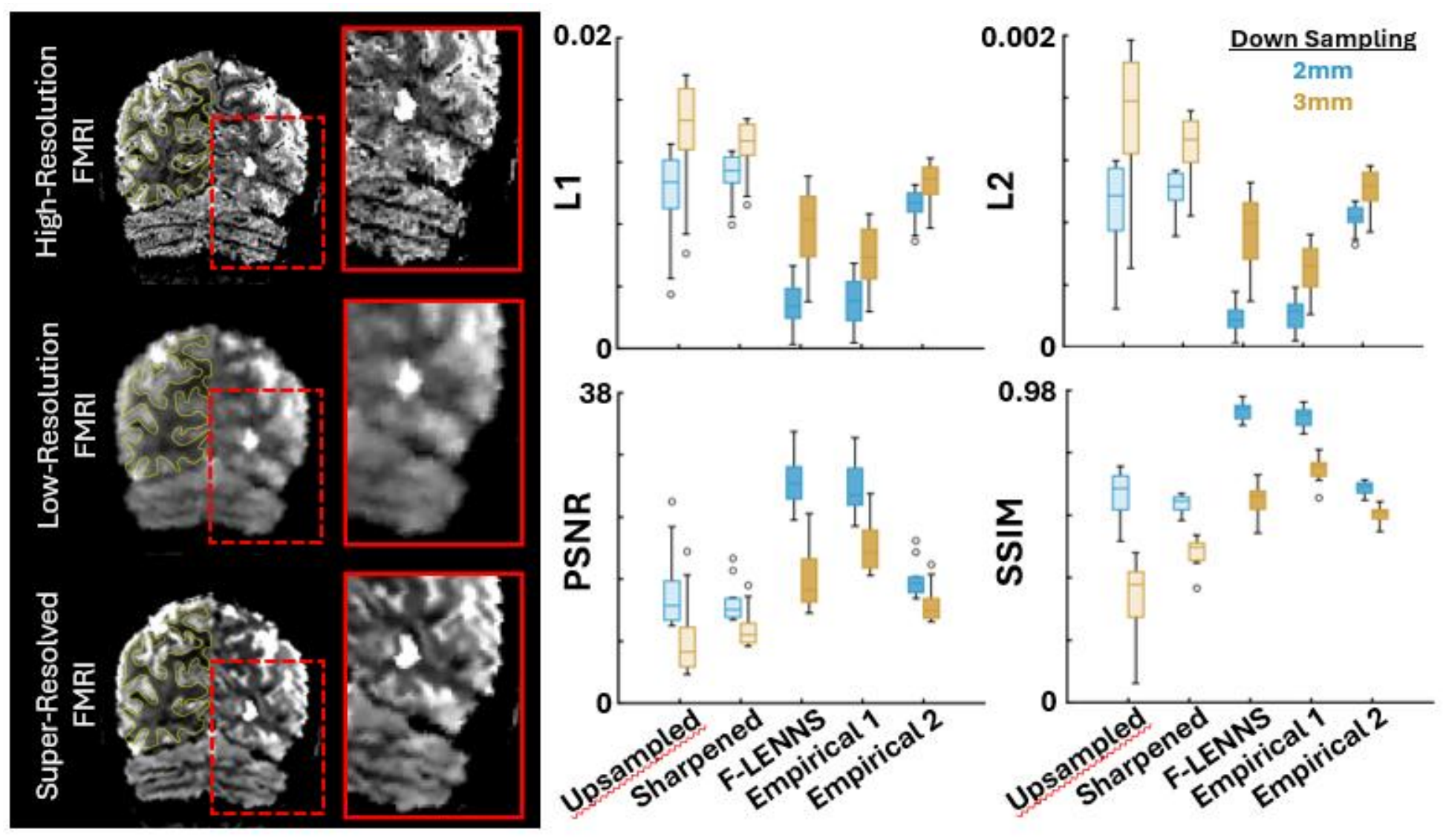
F-LENNS performance in improving the quality of raw fMRI images. The left panels show the high-resolution fMRI data (top), the corresponding blurred (downsampled) images (middle), and the super-resolved images generated by F-LENNS (bottom). The right panels show quantitative image quality metrics (L1, L2, PSNR, and SSIM) following upsampling, image sharpening, F-LENNS, and two other SR methods trained on real fMRI data. Empirical 1 represents a smaller domain gap, whereas Empirical 2 represents a larger domain gap.

It could be argued that training a model on empirical (real) rather than synthetic data might yield superior performance compared to f-LENNS. To test this, we trained two additional models with architectures identical to f-LENNS using empirical datasets. The first model was trained on data from six individuals acquired with identical scan parameters (i.e., minimal domain gap; see Methods), whereas the second model was trained on data from 90 individuals acquired with differing scan parameters, representing a larger domain gap. Individuals included in the validation set were excluded from both training sets.

When applied to 2 mm data, f-LENNS achieved performance comparable to that of the empirically trained model with minimal domain gap (t(10)=2.82; *p*=0.07) and substantially outperformed the model trained under a larger domain mismatch (t(10)=41.94; *p*<10⁻³). At 3 mm resolution (a much more challenging scenario where voxels are 27 rather than 8 times bigger), f-LENNS likewise remained superior to the more strongly mismatched empirical model, although a model trained under closely matched conditions achieved higher performance (t(10)=6.78; *p*<10⁻³). Together, these findings indicate that highly acquisition-matched empirical training can provide an additional performance advantage for particularly demanding SR tasks, whereas f-LENNS maintains strong performance across resolutions and domain shifts without requiring condition-specific training or model selection. Overall, f-LENNS provided robust image-quality enhancement despite domain mismatch, outperforming both AFNI and models trained on real datasets when the latter were applied across larger domain gaps.

### 2.3. F-LENNS enhances localizing mesoscale functional sites

We next evaluated whether these improvements in image appearance translated to gains in the downstream functional analyses one ultimately cares about. To test this possibility, we focused on the mesoscale organization of the occipital visual cortex, one of the most studied parts of the brain in humans^39,40^ and non-human primates^41–44^, and independently localized color-, motion-, and disparity-selective cortical sites in this brain region in eleven individuals with intact visual functions.

As demonstrated in Fig. 3, functional activity maps derived from the super-resolved data showed markedly improved localization of mesoscale structures within the occipital cortex, including motion- (Fig. 3A), disparity- (Fig. 3B), and color-selective (Fig. 3C) activity patches across early visual areas (V2-V4)^39,40^. Importantly, the relative location of these patches remains unchanged between the original and super-resolved maps (Fig. 3D).

**Figure 3.**
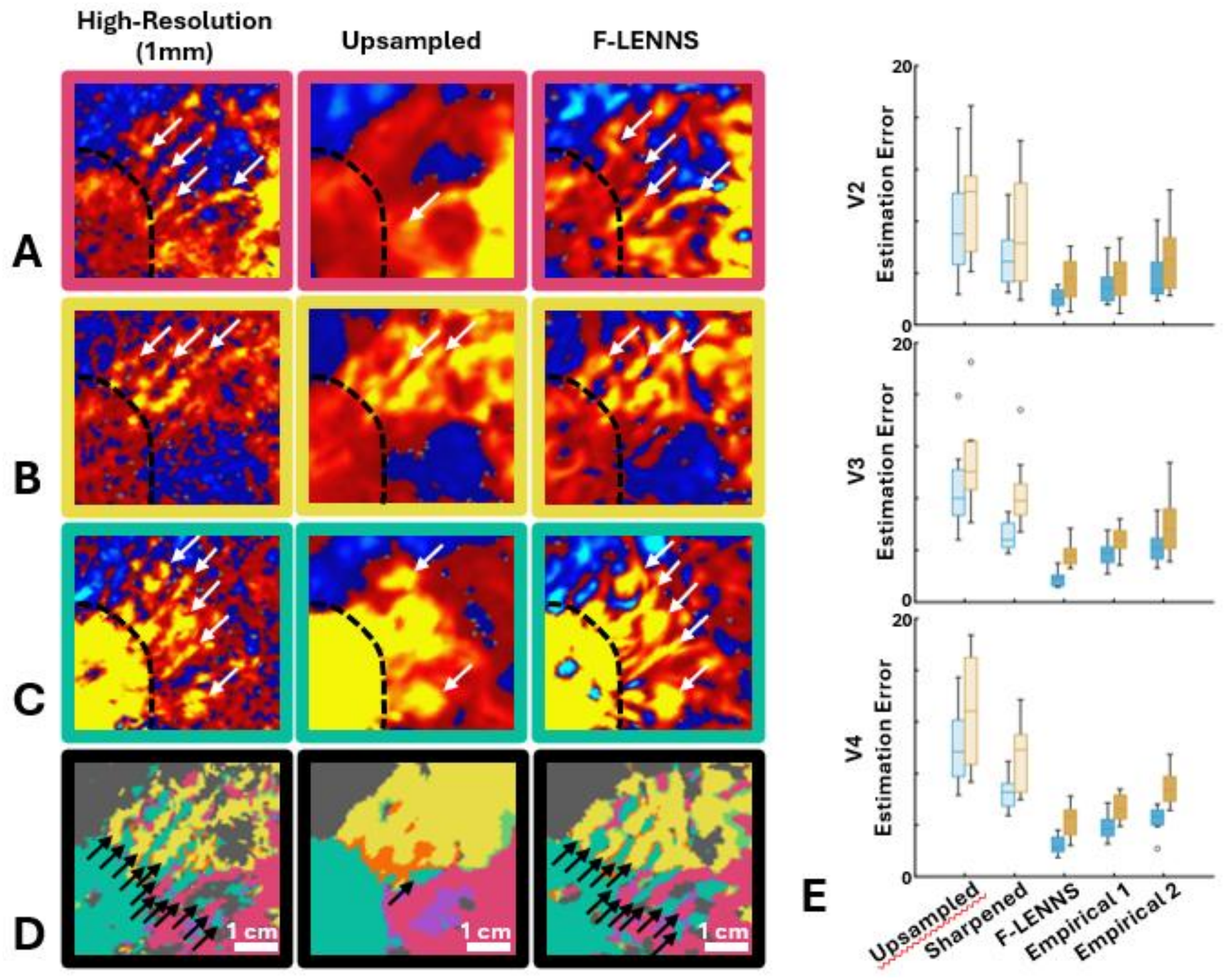
F-LENNS improves the localization of mesoscale functional activity patches across early extrastriate visual areas. Panels A–C show the unthresholded motion-, disparity-, and color-selectivity maps, respectively, from one representative participant across the early retinotopic visual areas. The border between striate (V1) and extrastriate (V2) visual areas is shown with dashed black line. Panel D shows the interdigitation of these functional patches after thresholding (for details, see ref. ^9,39^). In each panel, the left column shows the high-resolution data, the middle column shows the corresponding downsampled maps, and the right column shows the super-resolved maps generated by f-LENNS. Panel E shows the root-mean-square error (RMSE) in estimating the size of these functional patches across visual areas V2, V3, and V4 following upsampling, image sharpening, f-LENNS, and two SR methods trained on real fMRI data. Other details are the same as in Figure 2.

Quantitatively, for both 2 and 3 mm data, the estimated size of color-, motion- and disparity-selective activity sites in areas V2-V4, derived from the super-resolved data were significantly closer to those measured from the original high-resolution acquisitions than estimates obtained by upsampling the images and sharpening them with a classical image processing technique (unsharp masking) (t(10)>4.42; *p*<10⁻^2^) (Fig. 2E). Moreover, across V2-V4 and irrespective of the input data resolution, this estimate was comparable to (or slightly better than) those from the model trained based on the empirical data with minimal domain gap (t(10)>2.14; *p*<0.10) and superior to those obtained from the model based on the empirical data with larger domain gap (t(10)>2.75; *p*<0.02). Thus, our approach not only improved the raw image quality, but, crucially, it also enhanced the functional localization of mesoscale cortical structures and amplified their differential fMRI selective response.

### 2.4. F-LENNS generalizes to clinical populations

A key challenge for SR of fMRI is its validation in clinically relevant populations, where atypical functional organization raises the risk that the model could impose a normative template onto the data. To address this, we applied f-LENNS, along with the model trained with the empirical dataset with minimal domain gap, to fMRI datasets acquired from ten individuals with amblyopia, a condition known to selectively impair disparity processing while largely preserving color- and motion-selective responses.

As shown in Fig. 4, super-resolved fMRI data from amblyopic participants exhibited markedly reduced disparity-selective activity, consistent with their corresponding high-resolution data^9^, whereas color- and motion-selective maps remained comparable to those observed in neurotypical controls. Critically, these patterns were preserved after SR based on f-LENNS and the alternative model trained based on the empirical data with small domain gap, indicating that the model did not enforce typical functional organization to an atypically organized visual cortex. Moreover, patch size estimates derived from the super-resolved data were significantly closer to those measured from the original high-resolution acquisitions than estimates obtained from upsampled data, especially when the data is collected at lower spatial resolutions that is common in clinical studies (e.g., 3 mm; *p*<10^-3^). Thus, the super-resolved data faithfully reflected condition-specific alterations in functional responses, supporting the validity of the approach for studying individuals with atypical brain function.

**Figure 4.**
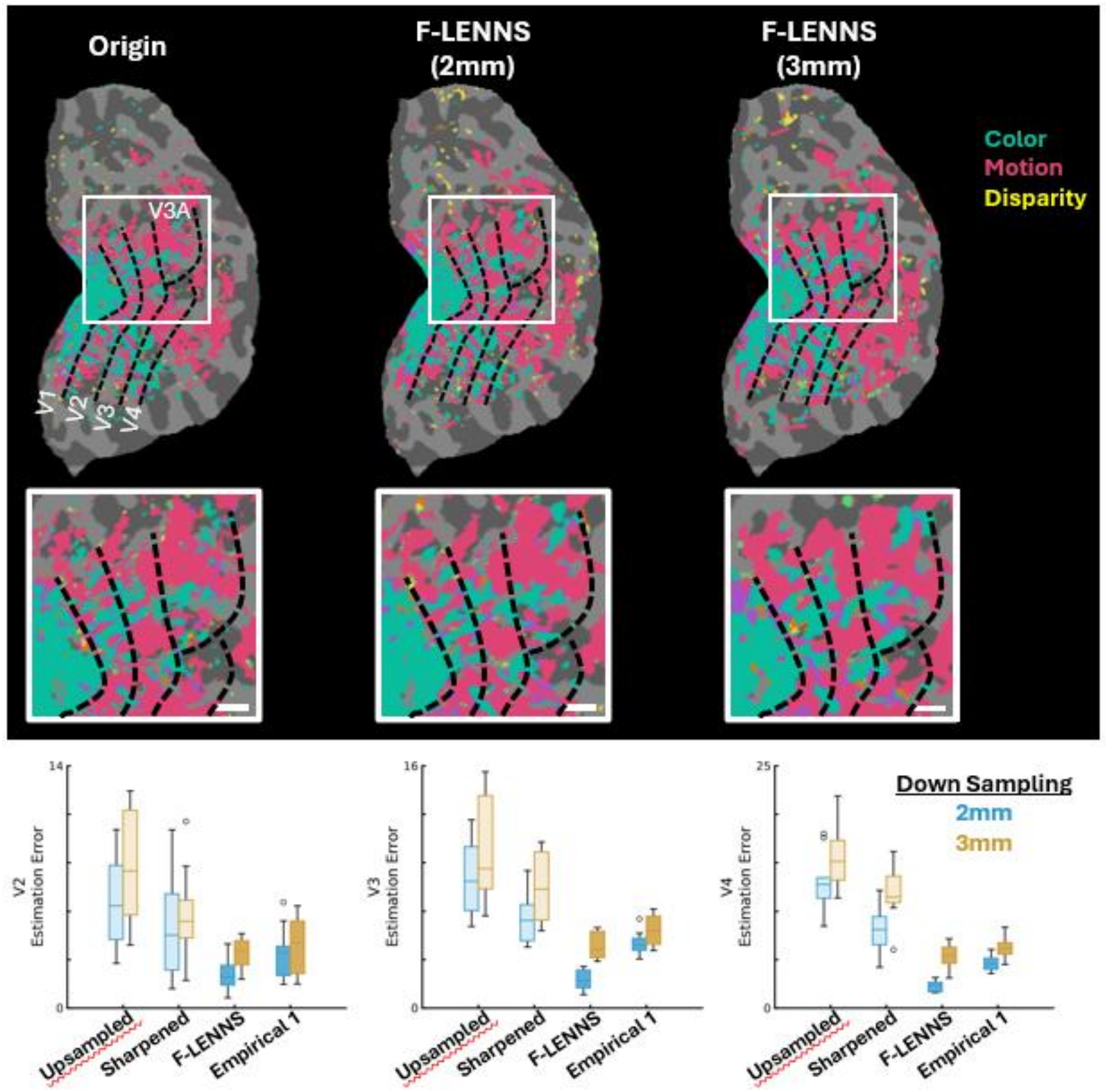
Application of f-LENNS to data from participants with amblyopia. The top panels show the interdigitation of color- and motion-selective sites and the absence of disparity-selective areas; this phenomenon is observed in disrupted binocular vision caused by amblyopia. From left to right, the panels show the original high-resolution data and the super-resolved reconstructions after downsampling to 2 mm and 3 mm resolution. The middle panels show magnified views of the inset regions from the top panels. The bottom row shows the error in estimating the size of these mesoscale sites using conventional upsampling, image sharpening, f-LENNS, and the proposed SR method trained on real fMRI data.

### 2.5. F-LENNS is applicable across field strengths

We next tested whether the same f-LENNS model could enhance fMRI data acquired at different magnetic field strengths by applying it to 3T data from participants who had also undergone high-resolution 7T imaging. Because f-LENNS is trained exclusively on synthetic images with randomized contrast and resolution, neither field strength constitutes a training domain in the conventional sense. Nevertheless, 3T and 7T fMRI differ in several properties that may influence the spatial organization apparent in the measured signal, including acquisition parameters such as TR and TE, signal-to-noise ratio, and the relative contributions of intravascular, venular, and extravascular components to the BOLD response. We therefore asked whether f-LENNS could enhance fine-scale functional organization in 3T data despite these acquisition- and field-strength-dependent differences.

To evaluate model generalization under these conditions, six participants were scanned at 3T in a separate session, using the same experimental paradigm as in the 7T acquisitions to localize their color-selective areas. As shown in Fig. 5, super-resolved 3T data revealed more pronounced mesoscale functional patterns across early visual areas compared to the upsampled and sharpened versions of the low-resolution data. Consistently, compared to simply upsampled and sharpened maps, the estimated number of cortical sites across visual areas V2-V4 based on f-LENNS application was closer to the number of sites detected in the high-resolution functional map from the same individuals collected in 7T (*p*<0.05).

**Figure 5.**
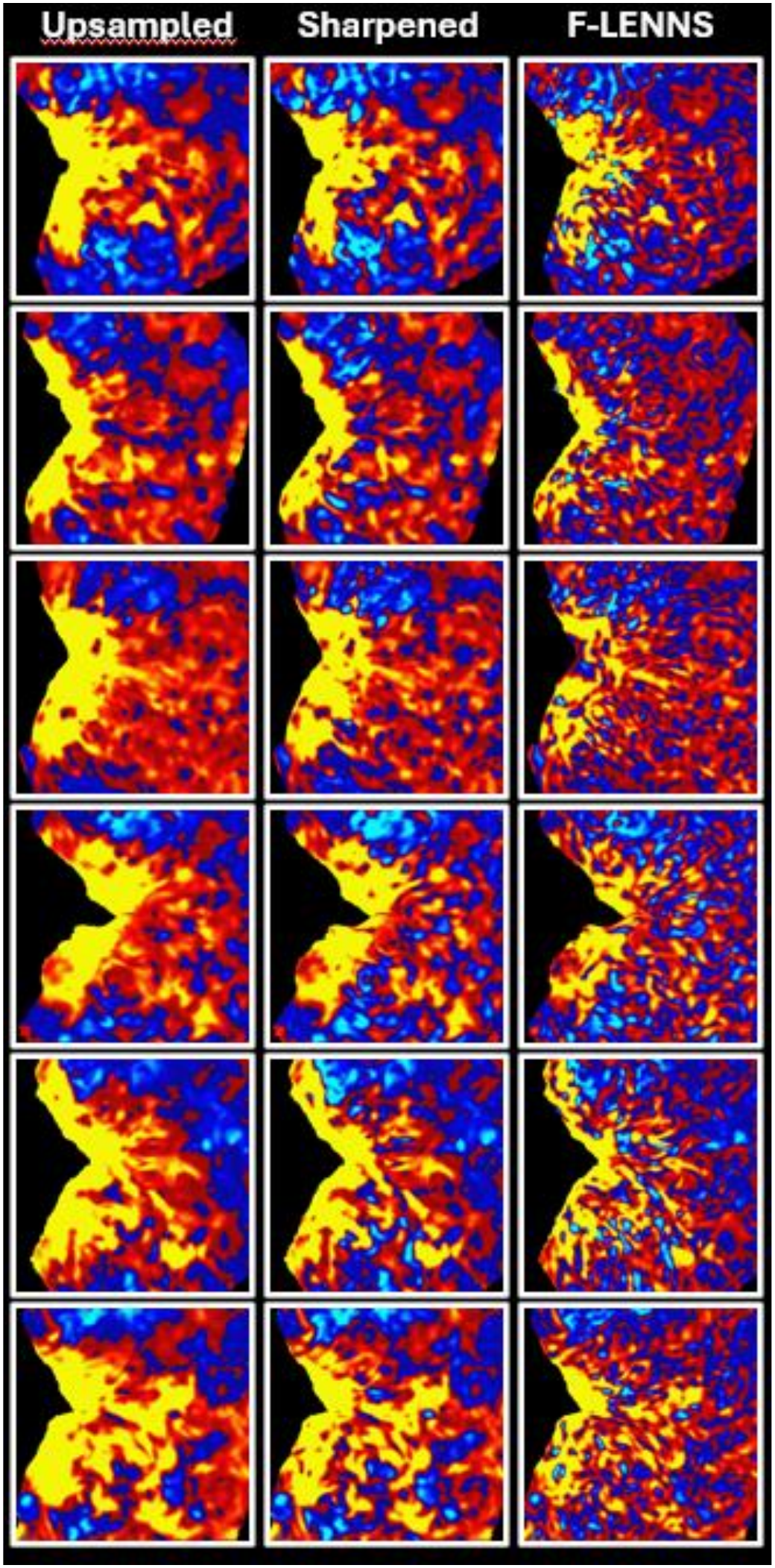
Application of f-LENNS to color-selectivity maps acquired from six participants on a 3T MRI scanner. Each row corresponds to one participant. The left column shows the original maps reconstructed at 3 mm spatial resolution (see Methods). The middle column shows the results after AFNI image sharpening. The right column shows the f-LENNS reconstruction. f-LENNS successfully recovered the characteristic stripe pattern across V2.

Importantly, even though the 3T data suffers from a much lower signal-to-noise ratio compared to 7T data, the 3T acquisitions can still provide a substantially larger field-of-view coverage, encompassing the whole brain. This result demonstrates that the proposed framework can enhance mesoscale functional organization in lower-field, whole-brain fMRI data, bridging differences in field strength and acquisition parameters.

## 3. Discussion

There is a growing demand for high-resolution fMRI to investigate mesoscale functional organization in the human brain, yet access to these approaches remains largely restricted by the availability of UHF MRI systems. Even when UHF scanners are available, acquiring high-resolution fMRI often requires compromises in temporal resolution, spatial coverage, or scan duration to maintain adequate signal- and contrast-to-noise ratios. Our results demonstrate that f-LENNS substantially alleviates these constraints by recovering fine-scale functional organization from conventionally acquired fMRI data while preserving biologically meaningful spatial structure. This approach is specifically useful to revisit publicly available datasets that were acquired using lower spatial resolution in the past, such as the Human Connectome Project (HCP)^45^. In doing so, it expands access to precision functional neuroimaging and provides greater flexibility in designing acquisition protocols at both conventional and UHF strengths. The same reasoning applies at the opposite end of the resolution range: the current ceiling for human fMRI, approximately 0.3 mm isotropic^46^, is imposed by MRI hardware rather than by the spatial scale of the underlying physiology. Because f-LENNS learns a mapping between resolutions rather than a dataset-specific prior, retargeting it to finer synthetic supervision offers a route to functional maps at resolutions currently unattainable *in vivo*.

### 3.1. Out-of-the-box resolution enhancement with broad generalizability and limited hallucination risk

One of the principal findings of this work is that robust SR of fMRI does not require dataset-specific training. Although models trained on matched high-resolution fMRI data can potentially achieve excellent performance when the training and test data closely resemble one another, their practical utility is inherently limited by the need to acquire matched high-resolution datasets for every imaging protocol. Such datasets are costly to collect and often impractical across institutions, scanner platforms, and clinical populations. Furthermore, adapting these models to new datasets typically requires repeated retraining, computational resources, and machine-learning expertise.

In contrast, f-LENNS achieves performance that approaches that of closely matched supervised models while maintaining high performance across a broad range of acquisition conditions using a single pretrained neural network. This robustness arises from training exclusively on domain-randomized synthetic data, enabling the model to generalize across varying field strength, pulse sequence, spatial resolution, scanner hardware, and experimental paradigm. As a result, f-LENNS can be deployed immediately, without retraining, fine-tuning, or parameter optimization, substantially lowering the barriers to adopting AI-based SR in functional neuroimaging. Furthermore, we have shown that domain randomization approaches the performance of fully supervised methods when training and test data are matched, and surpasses them (including narrower, more accurate simulations) when a domain gap (even if small) does exist.

Another benefit of training with domain-randomized Gaussian data is that the risk of hallucinating anatomical features is minimal, since the network only sees Gaussian cartoon-like images during training. We further minimize the hallucination risk by explicitly avoiding generative and adversarial components that are frequently used in the medical 3D SR literature, which improve the realism of the outputs at the expense of increased hallucination risk^47^.

### 3.2. Regaining autonomy in experiment design via f-LENNS

Over the past few years, the use of large publicly available high-resolution neuroimaging datasets, such as the HCP^45^ and the Natural Scenes Dataset (NSD)^48^, have significantly broadened access to high-quality fMRI and accelerated research into mesoscale brain organization. However, these datasets were designed to address specific scientific questions and therefore employ fixed acquisition protocols, participant populations, and experimental paradigms. Consequently, they cannot replace the flexibility required to investigate new hypotheses or specialized clinical cohorts.

Rather than relying on existing high-resolution datasets, f-LENNS enables investigators to recover mesoscale functional organization from their own conventionally acquired data. This capability allows experiments to be tailored to specific scientific questions, participant populations, and task designs while remaining compatible with widely available MRI systems. By decoupling fine-scale functional mapping from the availability of UHF hardware and predefined public datasets, f-LENNS broadens participation in mesoscale neuroimaging and may accelerate discovery across both basic and clinical neuroscience.

### 3.3. Frame-wise processing preserves compatibility with fMRI analysis

A distinguishing feature of f-LENNS is that each fMRI volume is processed independently. Although incorporating temporal information into deep learning models could potentially improve raw image SR under specific acquisition conditions, it also introduces learned temporal dependencies whose form and magnitude are not known a priori and may vary across experimental paradigms. Unlike conventional linear temporal filtering, for which the induced autocorrelation structure can be explicitly characterized and incorporated into voxel-wise statistical models, nonlinear temporal processing by a neural network alters the temporal covariance structure in ways that are difficult to quantify or correct for, largely complicating conventional fMRI statistical inference.

Frame-wise processing therefore involves a deliberate tradeoff: it forgoes potentially useful temporal information in exchange for preserving compatibility with standard fMRI analysis and avoiding opaque temporal dependencies introduced by multi-frame neural networks. It also allows the same model to be applied across paradigms and repetition times without assumptions about temporal structure. This broad applicability is especially important for publicly distributed software, where the acquisition characteristics and experimental design of future input data cannot be anticipated. Together with the generalization afforded by domain-randomized training, frame-wise processing therefore helps ensure that f-LENNS remains applicable across a wide range of real-world datasets. This flexibility may be particularly valuable in clinical applications, where temporal signal characteristics can differ because of altered neuronal or neurovascular dynamics and where imposing learned temporal relationships could reduce model reliability.

### 3.4. Limitations

F-LENNS is currently developed and validated on datasets acquired from individuals with anatomically intact brains. While the underlying domain randomization is highly generalizable and, in principle, applicable to data from brains with structural abnormalities, its performance in such settings remains to be systematically evaluated. Future work will be required to establish robustness and identify any necessary adaptations for these conditions.

In addition, f-LENNS is not designed to fully resolve intrinsic differences in the spatial organization of functional maps arising from variations in scanner platforms, pulse sequences (e.g., BOLD versus VASO), or field strengths (e.g., 3 T versus 7 T). A model tailored to the signal properties of each acquisition could potentially capture these differences more precisely and achieve a higher performance ceiling. Such an approach, however, would require detailed knowledge of the acquisition protocol and the selection or training of a separate model for each setting. By contrast, f-LENNS provides a single, acquisition-agnostic framework that can be applied across heterogeneous imaging conditions without requiring protocol-specific model selection. Although this generality may come at the cost of some pulse-sequence- or field-strength-specific optimization, our results show that f-LENNS nevertheless substantially improves the recoverability of fine-scale functional organization and supports more informative analyses across a broad range of acquisition scenarios.

Finally, although the present study focuses on visual cortex because its well-characterized mesoscale architecture provides a rigorous benchmark for quantitative evaluation, the underlying framework is not specific to the visual system. Future work will be needed to evaluate performance across additional brain regions and cognitive domains with distinct functional architectures.

### 3.5. Conclusion

We present f-LENNS, an AI-based framework that substantially enhances the effective spatial resolution of functional MRI while generalizing across heterogeneous acquisition protocols using a single pretrained model. Unlike existing approaches, f-LENNS operates “out of the box”, requiring neither retraining nor protocol-specific adaptation before application to new datasets. This combination of robustness and ease of deployment makes precision functional neuroimaging accessible to a substantially broader research community.

Beyond improving image quality, f-LENNS changes how high-resolution fMRI can be used in practice. By enabling retrospective enhancement of existing datasets and supporting prospective studies on widely available MRI systems, it expands opportunities for investigating mesoscale functional organization without requiring specialized hardware or acquisition-specific AI models. More broadly, this framework illustrates how domain-randomized learning can overcome longstanding challenges in cross-protocol generalization and may provide a foundation for the next generation of scalable, AI-assisted functional neuroimaging.

An exciting avenue for future work is to leverage emerging ultra-high-resolution anatomical datasets, such as ex vivo MRI and hierarchical phase-contrast tomography (HiP-CT^49^), together with advanced segmentation frameworks such as NextBrain^50^, to generate anatomically informed training data for super-resolution at submillimeter scales (e.g., ∼250 μm isotropic). While such resolutions are unlikely to be achievable through direct super-resolution of conventional 3-mm fMRI, ongoing advances in high-resolution functional imaging^51^ could enable progressively finer super-resolution, potentially extending the proposed framework well beyond the 1-mm resolution explored here.

## 4. Methods

### 4.1. Participants

Twenty-seven human adults (12 females), aged 19–56 years, participated in this study. Of these 27, 17 had normal or corrected-to-normal visual acuity and 10 had amblyopia (5 with anisometropic and 5 with strabismic amblyopia). Outside the scanner, the stereoacuity was measured for individuals with amblyopia using the Randot stereo test (Stereo Optical) to ensure their absence of measurable stereoacuity^9^.

All experimental procedures conformed to NIH guidelines and were approved by Massachusetts General Hospital protocols. Written informed consent was obtained from all participants prior to all experiments. The study conformed to the Declaration of Helsinki and was approved by the Massachusetts General Brigham Institutional Review Board.

### 4.2. Data acquisition procedure

All participants with neurotypical vision and no history of developmental and/or neuropsychological disorders were scanned in an UHF 7T scanner (whole-body system, Siemens Healthcare, Erlangen, Germany) for the functional experiments. Among them, six participants with neurotypical vision, who had more limited availability, were scanned only once during resting-state. These data was used for training the competing model based on real images.

The other eleven participants with neurotypical vision, along with the amblyopic individuals, were scanned multiple times in the 7T scanner to localize their motion-, disparity-, and color-selective sites, and to map their visual areas retinotopically.

All participants were also scanned in a 3T scanner (Tim Trio, Siemens Healthcare) for structural imaging. Six individuals with neurotypical vision, selected based on their availability, were also scanned to (re-)localize their motion-selective sites based on 3T scans, even though the rest of the procedure remained identical to that used in 7T scans (see below).

### 4.3. Stimuli

Details of stimuli are described previously ^9,39,40,52,53^. Briefly, disparity-selective sites were localized by comparing responses to disparity-varying stimuli (horizontal disparity: −0.22° to 0.22°, i.e., between ±792 arcsec; oscillated at 0.3 Hz) versus translationally moving, zero-disparity stimuli (left-to-right displacement: −0.22° to 0.22°; oscillates at 0.3 Hz)^9,40^.

Motion-selective columns were localized based on their stronger response to moving stimuli compared to stationary stimuli^39,52^. Stimuli consisted of concentric dark gray (30 cd/m^2^) rings, which were separated by 2° and had a width of 0.4°, extending over 20° × 26° (height × width) in the visual field, and were presented against a light gray background (40 cd/m^2^). In half of the blocks, rings moved radially (centrifugally vs. centripetally) at 4°/s, and the direction of motion changed every 3 s to reduce the impact of motion after-effects.

Color-selective sites were localized by presenting the participants with sinusoidal gratings (0.2 cycles/deg; 20° × 26°) that varied either in chromaticity (alternating between red and blue) or in achromatic luminance^40,53^. For each participant, red and blue hues were adjusted to be isoluminant across all stimulated eccentricities using the method of flicker photometry and tailored according to individual color perception^40,53^.

The border of retinotopic areas were defined by identifying representations of the vertical and horizontal meridians projected to surface reconstructions of the cortex^54,55^.

Resting-state data was collected during eye-closed conditions. Participants were instructed to stay still but not to go to sleep. Each scan took 264 s, and each participant was scanned 3-4 times. Experimenters talked to the subject between runs to ensure wakefulness.

### 4.4. Imaging

#### 4.4.1. Structural scans

We used anatomical images acquired at 3T on a Siemens TIM Trio whole-body scanner, equipped with the standard vendor-supplied 32-channel RF head coil array, which provides more uniform image contrast over the full field of view (FOV) than imaging at 7T. We used a 3D T1-weighted MPRAGE sequence with protocol parameter values: TR=2530 ms, TE=3.39 ms, TI=1100 ms, flip angle=7°, bandwidth=200 Hz/pix, echo spacing=8.2 ms, voxel size=1.0×1.0×1.0 mm³, and FOV=256×256×170 mm³.

#### 4.4.2. Functional scans

##### 7T scans

We utilized a 7T whole-body Siemens scanner equipped with SC72 body gradients (70 mT/m gradient strength and max slew rate of 200 T/m/s), a custom-built 32-channel helmet radiofrequency (RF) receive coil array, and a birdcage volume RF transmit coil. Functional data were collected using single-shot 2D gradient-echo EPI, with 1.0 mm isotropic nominal voxel size, and the following protocol parameter values: TR=3000 ms, TE=28 ms, excitation flip angle=78°, FOV matrix=192×192, BW=1184 Hz/pixel, echo-spacing=1 ms, 7/8 phase partial Fourier, 44 oblique-coronal slices covering the occipital lobe, acceleration factor R=4 with GRAPPA reconstruction and FLEET-ACS data^56^ with a 10° flip angle.

##### 3T scans

We utilized a 3T Tim-Trio Siemens scanner equipped with a 32-channel helmet RF receive coil array. Functional data were collected using single-shot 2D gradient-echo EPI, with 3.0 mm isotropic nominal voxel size, and the following protocol parameter values: TR=2000 ms, TE=25 ms, excitation flip angle=90°, FOV matrix=64×64, BW=2298 Hz/pixel, echo-spacing=1 ms, no phase partial Fourier, 33 Transverse slices covering the whole brain.

### 4.5. Data analysis

All real fMRI data used in this study, including the datasets used for evaluating f-LENNS and training the competing models, consisted of scanner-reconstructed magnitude images. Neither complex-valued images nor raw k-space data were used. Functional and anatomical MRI data were pre-processed and analyzed using FreeSurfer (version 7.4.0; http://surfer.nmr.mgh.harvard.edu)^57^ and in-house developed MATLAB code as described in detail in our previous publications^9,39,58^.

### 4.6. Machine learning model and training

#### 4.6.1. Architecture

The f-LENNS backbone is a volumetric adaptation of Nonlinear Activation Free Network (NAFNet)^36^ in which all operations are extended from 2D to 3D while preserving the model’s lightweight design. The network follows a U-shaped encoder-decoder topology with skip connections, composed of stacks of NAF blocks that replace conventional nonlinearities with channel-wise gating (i.e., element-wise multiplications) and avoid attention mechanisms to reduce memory and computational overhead. The model operates with a base width of 128 feature channels. The encoder comprises three stages operating at spatial scales of 1, 2, and 4 mm, defined relative to the common 1-mm isotropic input grid, with 2, 2, and 56 NAF blocks, progressively reducing spatial resolution via strided convolutions while increasing the receptive field. A bottleneck middle stage of 2 additional NAF blocks processes the lowest-resolution representation. The decoder mirrors the encoder with 2 blocks per stage and restores spatial resolution using sub-pixel convolution (PixelShuffle^59^ extended to 3D), followed by convolutional refinement and fusion with corresponding encoder features through skip connections. This parameter-free upsampling strategy is consistent with the original NAFNet design and avoids artifacts and overhead associated with transposed convolutions. Also consistent with NAFNet design principles, each block is built from lightweight residual units^60^ with simplified normalization and no explicit activation functions, which both stabilizes training and minimizes GPU memory usage (critical for 3D volumes). Overall, this configuration balances depth and efficiency, allowing high-capacity modeling of volumetric structure while remaining tractable for whole-brain inputs.

#### 4.6.2. Training data

Our neural network is trained on synthetic data produced by a generative model that requires 3D segmentations to guide the synthesis process. These 3D segmentations were obtained by automatically segmenting real 1×1×1 mm T1-weighted scans from 10 public brain MRI repositories spanning healthy aging and a broad range of neurological and psychiatric disorders (including autism, ADHD, Alzheimer’s, schizophrenia, and multiple sclerosis): ABIDE^61^, ADHD200^62^, ADNI^63^, AIBL^64^, COBRE^65^, Chinese-HCP^66^, HCP^67^, ISBI2015^68^, MCIC^69^, and OASIS3^70^). These scans were segmented with FreeSurfer^71^ and SAMSEG^72^ to obtain labels for cerebral and extracerebral structures, respectively. Visual quality control of these segmentations yielded a training dataset with 5,241 total subjects. The real T1 scans were immediately disregarded and are not used for training.

#### 4.6.3. Data generation and domain randomization

Training data for f-LENNS are entirely synthetically generated using a domain randomization framework^34^ designed to simulate extremely wide variability in brain MRI appearance while ensuring access to high-resolution supervision. Realistic neuroanatomical structure is provided by 3D segmentations from the 5,241 subjects in the training data. Given these anatomical labels, synthetic imaging volumes are generated via a probabilistic forward model combining geometric augmentation (affine and nonlinear), Gaussian mixture tissue intensity modeling, and spatially smooth bias field perturbations^37,73^. Importantly, these synthetic volumes are scalar, magnitude-like intensity images (i.e., Gaussian “cartoons”), rather than simulations of complex-valued MR signals. The Gaussian distributions model image intensities directly; no phase information is generated or used during training.

This generative process enables explicit control over tissue contrast, intensity distributions, and scanner-induced inhomogeneities. To further enhance robustness, acquisition resolution is randomized at each training iteration, with voxel sizes sampled from a distribution specifically tailored to the range of spatial resolutions commonly encountered in fMRI acquisitions (uniform between 2 and 4.5 mm). Low-resolution inputs are simulated by resampling the high-resolution synthetic images to arbitrary voxel spacings and then interpolating them back to a common 1 mm isotropic grid. The network receives only these interpolated images, without any explicit information about the original voxel size used to generate them. It is therefore trained to recover high-resolution image details across a range of input resolutions while preserving the original image contrast, without being explicitly conditioned on voxel sizes.

This formulation implements domain randomization at both the anatomical and acquisition levels, exposing the network to a wide range of plausible imaging conditions. This fully synthetic and highly variable training regime allows the model to learn a mapping that is invariant to scanner-specific characteristics and input resolution, enabling inference on heterogeneous data without retraining or fine-tuning.

#### 4.6.4. Real data for training of competing models

We used two different datasets with real fMRI data to assess the impact of training on empirical versus domain-randomized data. The first dataset comprises resting-state fMRI data from six participants, none of whom were included in the f-LENNS evaluation dataset. The acquisition parameters^74^ were almost identical to those used in the f-LENNS evaluation, ensuring consistency between the training and evaluation datasets. During scanning, participants were instructed to keep their eyes closed, remain awake, and avoid falling asleep.

The second real dataset, which we used to assess the impact of the domain gap, is an independent resting-state fMRI dataset that is publicly available^75^. It comprises 90 participants who underwent a 10-minute resting-state fMRI scan on a 7T MRI scanner, while keeping their eyes open. Scanning parameters (including TR, TE, flip angle, and FOV matrix size) differed substantially from those used for evaluating f-LENNS, thus creating a larger domain gap.

#### 4.6.5. Training procedure

Our model was trained to perform contrast-preserving voxel-wise SR by minimizing the mean absolute error (L1 norm) between predicted and reference 1×1×1 mm synthetic images. This objective encourages the recovery of fine spatial details while maintaining the original image contrast, an important property for enhancing fMRI data acquired using different pulse sequences and contrast mechanisms.

The L1 norm encourages accurate intensity reconstruction while being more robust to outliers than the L2 norm (root of sum of squared differences). We do not employ perceptual losses based on deep generative models, as these can introduce hallucinated high-frequency content in pixel-level regression tasks^76^ and are therefore undesirable for anatomically faithful reconstruction. Optimization was performed using the Adam optimizer^77^ with a learning rate of 0.0001 and default momentum parameters, providing stable convergence under the high variability induced by domain randomization. Training proceeds on fully synthetic data with randomized resolution and appearance at every iteration, which effectively eliminates the need for a separate validation set. Instead, convergence is assessed empirically by monitoring the stabilization and plateauing of the training loss over a large, fixed number of iterations (100,000).

#### 4.6.6. Other implementation details

Due to memory constraints associated with 3D volumetric processing and to preserve computational frugality, training is conducted on randomly sampled 64×64×64 voxel patches extracted from the synthetic volumes (batch size: 1). This provides a balance between local anatomical detail and computational efficiency while maintaining sufficient context for learning SR mappings. Input intensities are min–max normalized to the range [0,1] prior to being processed by the network, ensuring numerical stability and consistent dynamic range across synthetic samples. At inference time, real fMRI images are similarly resampled from their native voxel size to the common 1 mm isotropic grid before being passed to the network. The original voxel size is used only for this resampling step and is not provided to the network as an additional input. The model then operates on full 3D volumes without patch-wise decomposition. Inputs are first padded to the nearest multiple of 4 in each spatial dimension to ensure compatibility with the encoder–decoder downsampling and upsampling scheme. The entire volume is then processed in a single forward pass, leveraging the fully convolutional nature of the architecture and avoiding tile-based stitching artifacts. This design enables efficient deployment while preserving spatial consistency.

### 4.7. Statistical analysis

Paired t-test was used primarily to test the impact of the enhancement method. All tests were corrected for multiple comparisons using Bonferroni correction.

### 4.8. Data sharing statement

F-LENNS codes are available at: https://github.com/jeiglesias/f-LENNS.git

## Acknowledgment

This work was primarily supported by NIH grants R01 EY030434 and RF1 AG080371, with additional support from NIH grants R01 AG070988, R01 EB031114, R01 CA296008, UM1 MH130981, R21 NS138995, and U19 AG060909, as well as the MGH/HST Athinoula A. Martinos Center for Biomedical Imaging. Essential resources were provided through NIH Shared Instrumentation Grant S10 RR019371 and NIH grant P41 EB030006. MSR gratefully acknowledges the generous support of the Kiyomi and Ed Baird MGH Research Scholar Award.

